# Drought dynamics explain once in a century yellow fever virus outbreak in Brazil with implications for climate change

**DOI:** 10.1101/2025.02.25.640139

**Authors:** Jamie M. Caldwell, Bryan Grenfell, Gabriel Vecchi, Joelle I. Rosser

**Affiliations:** High Meadows Environmental Institute, Princeton University, Princeton, 08544, USA; Department of Ecology and Evolutionary Biology, Princeton University, Princeton, 08544, USA; Department of Geosciences, Princeton University, Princeton, 08544, USA; Stanford University School of Medicine, Stanford University, Stanford, 94305, USA

## Abstract

While excess rainfall is associated with mosquito-borne disease because it supports mosquito breeding, drought may also counterintuitively increase disease transmission by altering mosquito and host behavior. This phenomenon is important to understand because climate change is projected to increase both extreme rainfall and drought. In this study, we investigated the extent to which seasonally-driven mosquito and primate behavior drove the first yellow fever virus (YFV) epidemic in an urban area in Brazil in nearly a century, coinciding with an equally rare drought, and to assess the role of interventions in ending the outbreak. We hypothesized that drought triggered the outbreak by driving the forest mosquitoes and non-human primates towards the city in search of water and that the mosquitoes were biting more frequently to avoid desiccation. A dynamical YFV model supports these hypotheses, showing that both behavioral changes were needed to explain the outbreak timing and incidence. Fur-ther, a combination of vector control, conservation measures, and vaccination contributed to ending the outbreak, with the strongest effects from vaccination. Together, these results suggest that drought, likely to become more frequent in this region in the coming decades, can significantly influence mosquito-borne disease transmission, and that sustained control will require multiple interventions.

## 1 Introduction

After nearly a century without urban yellow fever virus (YFV) transmission, Brazil experienced a major outbreak in 2017-18 (*1,2*), likely driven by a combination of land use change, viral evolution, accumulation of susceptible populations, and favorable climate conditions. YFV causes disease in humans and non-human primates with clinical presentations ranging from asymptomatic to life-threatening. YFV spreads via an urban transmission cycle involving *Aedes aegypti* mosquitoes and humans, or a sylvatic (i.e., forest) cycle between forest mosquitoes, non-human primates, and occasionally humans. By the 1930s, aggressive vector control and vaccination had controlled the urban cycle (*3, 4*), with ongoing YFV transmission in Brazil occurring exclusively through the sylvatic cycle. In recent years however, intensive land use change facilitated increased transmission of YFV into rural human populations at the forest edge (*5*). As deforestation increased, YFV spread southeastward in Brazil leading up to the 2017-18 outbreak (*6*), which was first reported in Minas Gerais and then in neighboring states of Espiritu Santo, Rio, and Sao Paulo (*1, 7*). The virus may have evolved increased transmutability during this time through phenotypic changes in nonstructural proteins (*8*), and historically low vaccination rates due to low YFV risk (*6, 9, 10*) left a large pool of susceptible individuals despite the high efficacy of the vaccine (*11*). Additionally, abundances of the urban mosquito, *Aedes aegypti*, were heavily reduced at the start of the YFV epidemic due to extensive vector control efforts in response to the 2015-2016 Zika outbreaks (*7, 12*). Together, these factors suggest that sylvatic mosquito vectors were carrying more transmissible strains of the virus and living in closer proximity to dense human populations with limited vaccine coverage (*6*). Against this backdrop, an extreme drought (*7,13*) may have triggered the outbreak by increasing contact between infected mosquitoes and humans, consistent with models identifying low vaccine coverage, high population density, and climate factors as key drivers of YFV spillover risk (*14*).

Given the role of water in the mosquito life cycle, there is an intuitive reason to suspect excess precipitation to increase YFV incidence, however, drought could also influence disease dynamics by concentrating hosts and vectors in the same locations and times and/or by increasing mosquito feeding to counteract dehydration. The YFV epidemic peaked during the rainy seasons in 2017 and 2018; during the extreme drought (*7, 13*), the modest rainfall during those periods may have intensified feeding and contact: water-scarce conditions drive both mosquitoes and non-human primates to travel farther for water, with mosquitoes biting more to avoid desiccation. During such scarcity, highly habitat adaptable and YFV-susceptible non-human primates, particularly howler monkeys and marmosets, are likely to concentrate in high densities in urban and peri-urban areas (*15, 16, 17, 18*), where high infection rates were observed (*1*). Similarly, *Haemagogus* forest mosquitoes, capable of flying long distances in search of food and water (*19*) would similarly move towards urban areas and increase biting under dehydration stress (*20*). Indeed, multiple lines of evidence show that forest mosquitoes, primarily in the genus *Haemagogus*, were responsible for YFV transmission during this epidemic (*1, 21, 18, 16, 22*). This concentration of non-human primates and forest mosquitoes could therefore amplify transmission of YFV within their own populations and in closer proximity to humans (*7*). Drought-induced migration of non-human primates and forest mosquitoes into urban areas could therefore result in a major human outbreak. This hypothesis is consistent with other studies suggesting that changing environmental conditions and increased contact between non-human primates, forest mosquitoes, and humans propagated this unusual outbreak (*2, 22*). Despite the biological plausibility of extreme drought triggering this YFV outbreak and other examples of drought concentrating pathogens, animals, and humans to amplify infectious disease transmission, drought has not previously been implicated in YFV outbreaks.

To evaluate the extent to which seasonally-driven mosquito feeding and non-human primate incursion into the cities could have contributed to the 2017-18 outbreak and assess intervention strategies, we simulated YFV transmission with a compartmental model parameterized based on different hypotheses and compared those outputs with real world evidence. We developed a compartmental model that captures key features of the Brazil YFV epidemic, including transmission from *Haemagogus* and *Aedes aegypti* mosquitoes to non-human primates (howler monkeys and marmosets) and humans (Figure 1). Within this model structure, we asked how well the model reproduced disease dynamics in Minas Gerais during the epidemic with nested models accounting for seasonally-driven mosquito feeding behavior, movement of forest non-human primates and mosquitoes, or both. Then we assessed the extent to which different interventions could have reduced the second wave of transmission and if those interventions would carry into the future.

**Figure 1:**
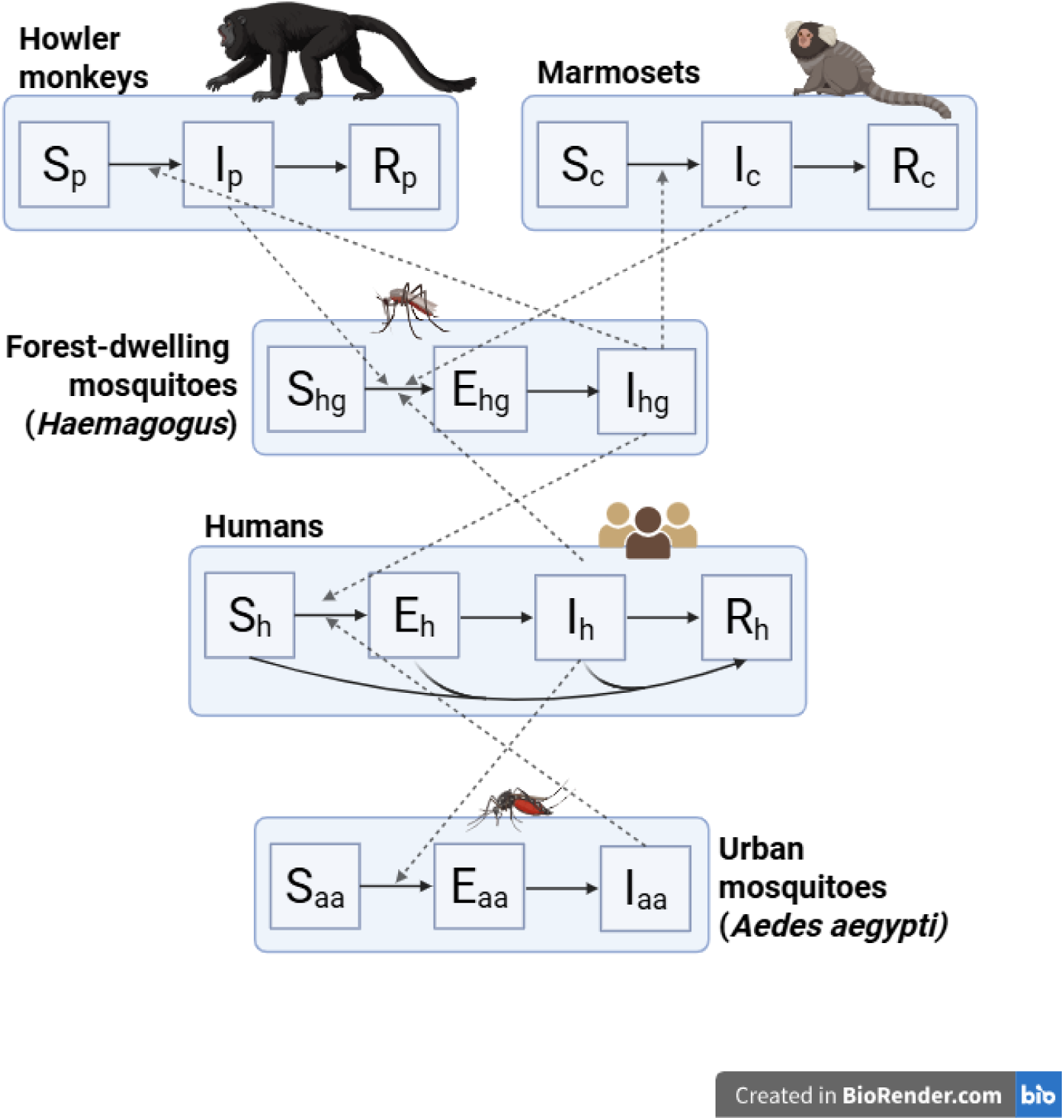
Compartmental model of yellow fever virus transmission in Brazil. Each blue box indicates a different species involved in disease transmission. Individuals within each species can move sequentially through several health states: susceptible (S), exposed (E), infectious (I), and removed (R). Non-human primates do not include an exposed state because it is unknown whether they have a latent period and how long it would last. Mosquitoes do not have a removed state because their lifespan is short relative to the study period. Solid arrows indicate the direction individuals move between compartments and dashed arrows indicate the direction of transmission. The curved arrows in the human population represent vaccination.

## 2 Results

### 2.1 Comparison of seasonal drivers

The model that incorporated both seasonally-driven mosquito biting behavior and movement of forest species (hereafter referred to as the *full model*) reproduced the observed two-year epidemic dynamics in both non-human primates and humans better than the nested versions of this model (Figure 2, Table 1). The full model captured the epidemic seasonality well and a sensitivity analysis supports the robustness of these conclusions (fig S1, table S2). If either or both seasonal components were excluded from the model, the predictions poorly predicted non-human primate epidemic dynamics and incorrectly reproduced the timing of the first epidemic wave (either too early or late) while failing to predict a second wave of transmission in humans. The nested models that incorporated seasonally-driven mosquito biting but restricted movement of forest species predicted no transmission in non-human primates and an early first wave of human transmission. Further, the model that included only seasonally-driven mosquito biting over-predicted human incidence in the first wave; it did however also predict a small increase in cases that did not materialize into an outbreak about four months before the second wave of transmission. In contrast, the model that incorporated seasonally-driven forest movement but restricted mosquito biting predicted a late first wave of human transmission; this model did show some uncertainty around the second wave of transmission at the correct time. These results indicate that both increased mosquito biting during the rainy season and increased movement of forest species into the cities, partially driven by the ongoing drought conditions, were needed for the two-peak epidemic to develop.

**Figure 2:**
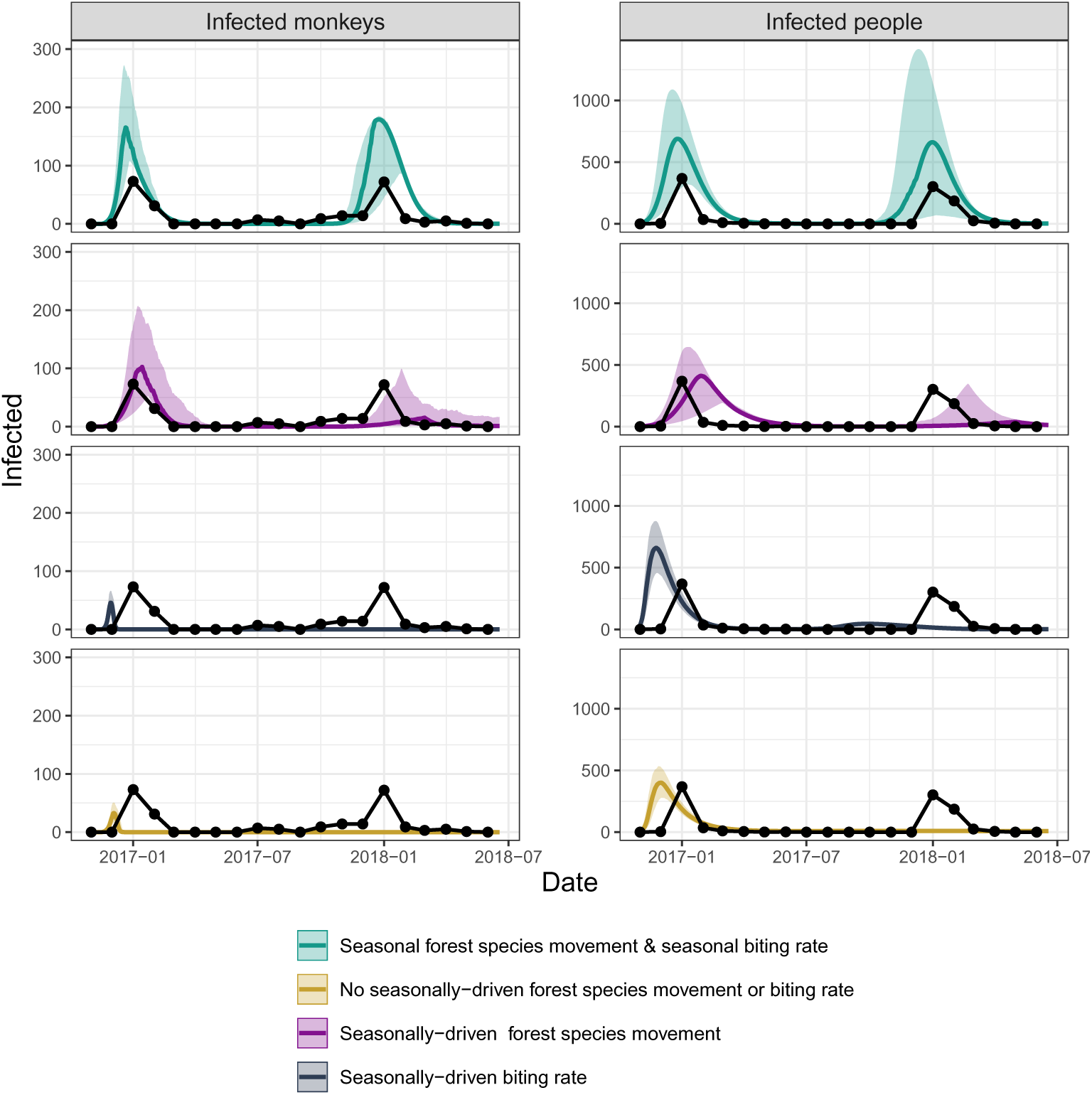
Seasonally-driven forest species movement and mosquito biting contributed to driving the two-wave YFV epidemic in Brazil. Comparison of four nested models with observations for non-human primate cases (left) and human cases (right, lead time = one month). The full model (teal) included seasonally-driven forest species movement and mosquito biting. The model with seasonally-driven mosquito biting (purple) holds forest species movement constant white the model with seasonally-driven forest species movement (dark blue) holds seasonally-driven mosquito biting constant; both models did not predict a second epidemic peak. The model that holds both seasonally-driven forest species movement and mosquito biting constant (yellow) predicted an early first wave and no second wave of transmission. We show observations overlaid for comparison (black dots connected by lines).

**Table 1:**
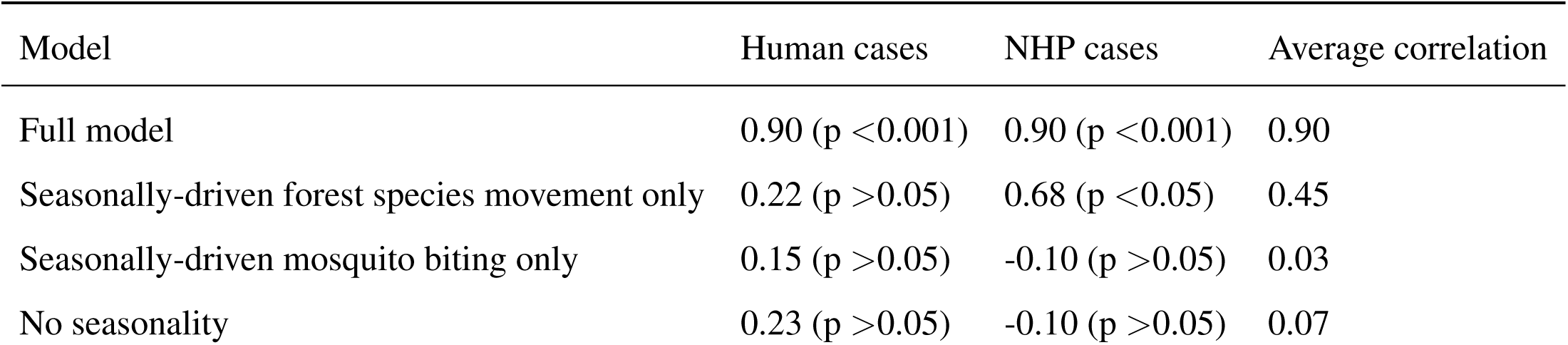
Correlation (p-value) between model predictions and observations across nested seasonally-driven models. Modeled human cases were adjusted by a one month lead time. NHP: non-human primates. Note that the’No seasonality’ model still includes a seasonal mosquito carrying capacity.

| Model | Human cases | NHP cases | Average correlation |
| --- | --- | --- | --- |
| Full model | 0.90 (p <0.001) | 0.90 (p <0.001) | 0.90 |
| Seasonally-driven forest species movement only | 0.22 (p >0.05) | 0.68 (p <0.05) | 0.45 |
| Seasonally-driven mosquito biting only | 0.15 (p >0.05) | -0.10 (p >0.05) | 0.03 |
| No seasonality | 0.23 (p >0.05) | -0.10 (p >0.05) | 0.07 |

### 2.2 Optimizing Interventions

Employing multiple interventions simultaneously had the highest likelihood of minimizing YFV transmission in the long term, but would not have been enough to prevent the second epidemic wave (Figure 3). The intervention that reflected spraying insecticides every rainy season, the key vector control tactic used in the region, led to a reduction in cases in the shorter term but allowed for increasingly large annual outbreaks due to reaccumulation of mosquito populations. The intervention that reflected conservation initiatives aimed at reducing movement of forest species to cities had a similar effect. The third intervention initiated the vaccination campaign at the onset of the epidemic (whereas in reality the campaign was delayed until after the first peak) and then continued YFV vaccination as part of routine childhood vaccinations in the region; this intervention had the strongest effect overall, with increasing effectiveness through time, contrasting the patterns of the first two interventions. The simulation that included all three interventions showed substantial and sustained control of YFV, which was largely driven by vaccination efforts. The other interventions still remain valuable though; if the virus evolved to evade the vaccine, or vaccine uptake or distribution were heterogeneous (as is often the case), the other interventions would provide a substantial buffer. In the hypothetical situation where the virus evolved to be 50% more transmissible, which would be possible in the future, we would expect much larger initial outbreaks and the need for stronger integrated control. Surprisingly, none of the interventions were strong enough to curb transmission in short term.

**Figure 3:**
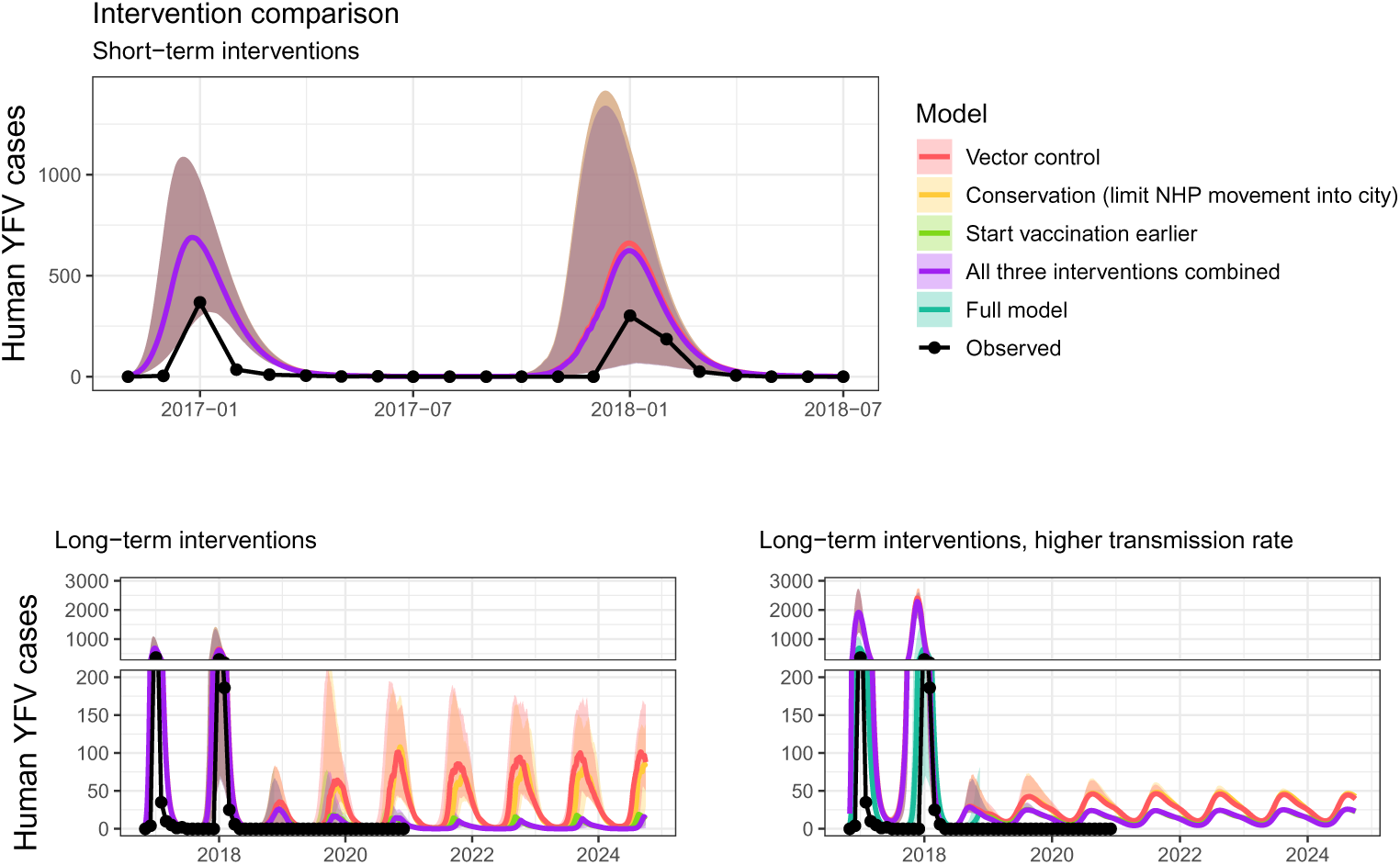
Deploying multiple control strategies simultaneously allow for sustained YFV control. Affects of interventions in the short-term (top) and long-term (bottom). We simulated four intervention strategies: vector control (i.e., reducing mosquito populations in the rainy season; pink), conservation initiatives (i.e., reducing forest species movement into the cities; orange), early and continued vaccine distribution (green), and a combination of all three (purple). For comparison, we show observations (black points connected by lines) and the full model (teal).

### 2.3 Extreme drought and global change

The drought prior to the 2017-18 YFV outbreak in Minas Gerais, Brazil was 9.5% more extreme than a once in a century drought (i.e., 100-yr return period) and dry conditions are projected to increase in the future (Figure 4). Based on the widely used drought metric, Standardized Precipitation Evapotranspiration Index (SPEI), we show that extreme dry conditions were present four months prior to the YFV outbreak. This timeline aligns with the expected bio-physical process that links climate to disease transmission whereby dry conditions promote migration of non-human primates and forest mosquitoes towards city centers in search of food and water in the months preceding the outbreak. Based on climate models, future conditions in this region are 47% drier than historical conditions at the 1% exceedance threshold, indicating a substantial increase in the severity of extreme drought events (based on projections with intermediate greenhouse gas emissions: SSP245). The IPCC-AR6 supports that drought should increase in Brazil (Figure 12.4 from (*23*)).

**Figure 4:**
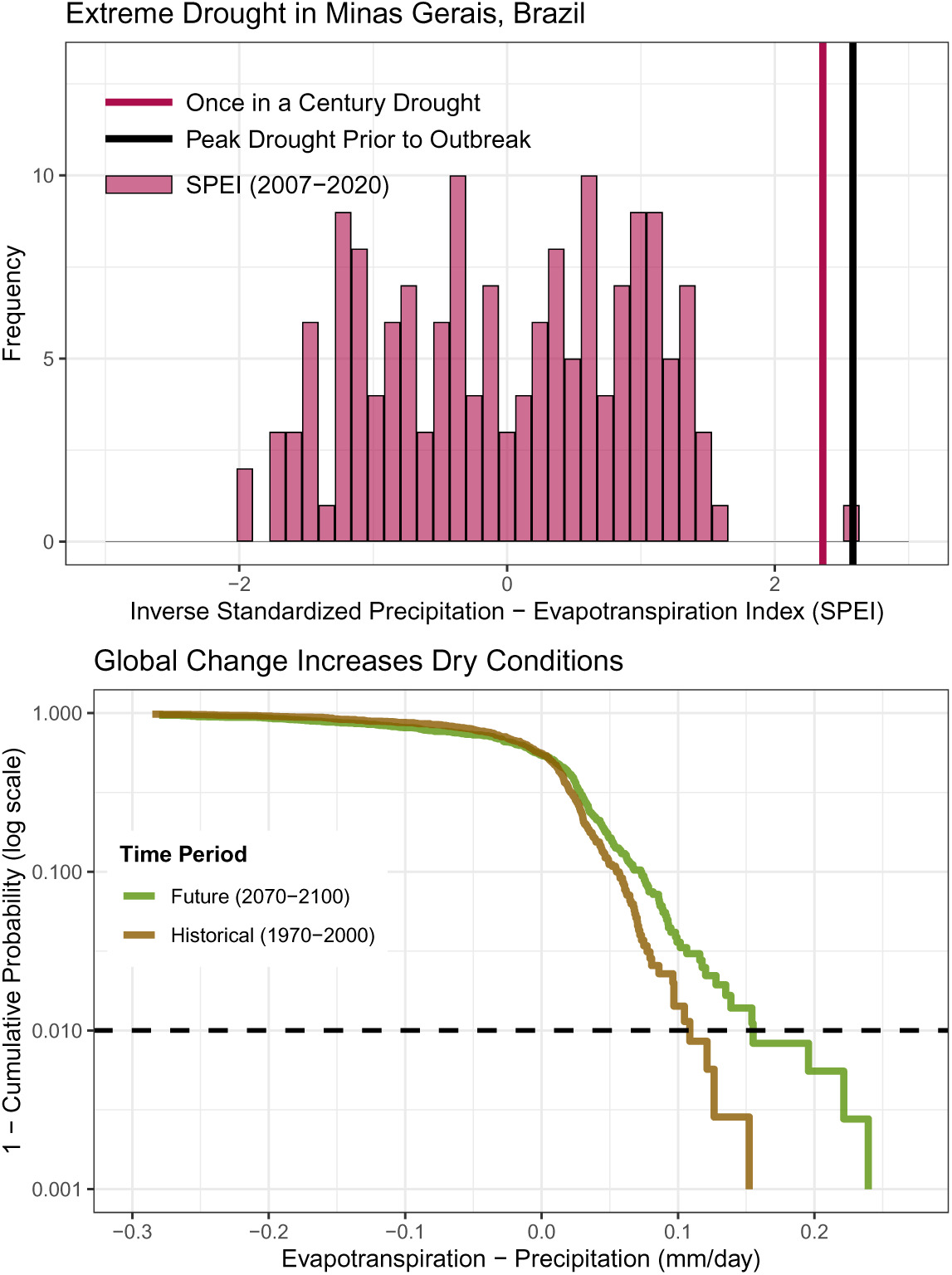
Conditions preceding the YFV outbreak were more extreme than a once in a century drought and global change will give rise to more extreme dry conditions in the future. Histogram of the inverse Standardized Precipitation-Evapotranspiration Index (SPEI) for Minas Gerais, Brazil (top). We show the value for a once in a century drought (pink vertical line) and the value for SPEI preceding the YFV outbreak (black vertical line). A cumulative distribution function (CDF) plot (bottom) shows the survival probability (1 - CDF) of the difference between evapotranspiration and precipitation (mm/day) for two time periods: Historical (1970–2000) and Future (2070–2100). We plotted the y-axis in log space to highlight extreme events, with a dashed horizontal line indicating a 1% exceedance threshold. The plot illustrates an increased likelihood of extreme dry conditions in the future compared to the historical period. For top and bottom plots, negative number indicate wetter conditions and positive numbers indicate drier conditions.

## 3 Discussion

Yellow fever virus (YFV) outbreaks are notoriously difficult to predict, and retrospective analyses can provide some insight into why outbreaks begin and why they end. The 2017-18 outbreak in southeastern Brazil was particularly unusual because there had not been transmission in the region in nearly a century. Prior studies hypothesized that slow changes (e.g., build up of a highly susceptible population, land use change) provided the background conditions for an outbreak to emerge (*14*), while we tested the hypothesis that seasonal climate conditions were responsible for the two-peak epidemic trajectory (*7*). By comparing models representing different hypotheses about seasonal drivers with real world observations, we show that the combined effects of seasonally-driven mosquito feeding behavior and seasonally-driven forest species incursion into the city explains the outbreak timing and incidence. Even without drought conditions however, it is possible that once seeded, an outbreak would have taken off, simply due to high susceptibility in the population. While seasonal climate appears to explain how the outbreak developed, it is equally important to understand why the outbreak did not continue to cause annual epidemics thereafter. The intervention simulations show that a combination of activities likely contained the outbreak, but could not have avoided the second peak in transmission altogether. In reality, the three techniques described (vector control, conservation, vaccination) were employed to different extents and largely align with real-life observations, where YFV cases were not reported after 2018. The simulations also suggest that there could be very low levels of continued transmission. Given the high rate of asymptomatic transmission (*24*), it is plausible that some cases go undetected every year.

The Brazilian epidemic was also unusual because it spread in urban areas but was predominantly spread by a non-urban mosquito species, amplified by non-human primate transmission. The outbreak was driven by *Haemagogus* rather than *Aedes aegypti* mosquitoes. The expectation was that given sufficient cases, there would be re-establishment of an urban transmission cycle perpetuated by *Aedes aegypti*–human transmission as *Aedes aegypti* remains a competent YFV vector in laboratory studies (*25, 26*). However, no infected *Aedes aegypti* were found during this outbreak (*27, 28, 29, 28, 30*). Thus, this outbreak was largely driven by *Haemagogus* flexibly feeding on alternate hosts, which has been reported previously (*31, 32, 33*), and was likely bolstered by feeding on non-human primates in the city, which have higher and longer viremia profiles than people (*1, 2*) indicating even low abundances of non-human primates can impact transmission. The contribution of non-human primates in urban transmission may also be the reason why vaccination alone did not appear adequate to reduce transmission in the short term. Our expectation was that to limit outbreaks (i.e., R_0_ *<*1), vaccination coverage needed to be 76-79% (coverage required to limit outbreaks = 1 - 1/R_0_ where median R_0_ = 4.21 and mean R_0_ = 4.81 across multiple studies (*34*)). In this study, we allowed vaccination coverage to reach 90%, yet the simulations showed vaccination alone could not have prevented the second wave of transmission, suggesting we should consider the susceptible population as a combination of people and non-human primates. A related important consideration is that continued viral evolution will increase the critical population size necessary for disease control. Thus, a suite of integrated control measures will become increasingly important.

Land use change was likely the proximate cause of non-human primate incursion into the city and ecological countermeasures may be key to reducing future disease emergence risk. Deforestation and other types of habitat alteration can increase spillover risk of disease from the forest to the city by affecting non-human primate susceptibility and spatial behavior (*35, 36*). Reduced habitat quality and area often limit food and water availability, forcing non-human primates to invest extra physiological resources for basic survival, leading to reduced immune function. Immune dysregulation can both increase susceptibility to infection and increase pathogen loads and shedding time (*35*). Additionally, animals may need to expand their search area to find food (*35, 37, 38, 39*). These behavioral changes then increase the likelihood and length of infected non-human primates-*Haemagogus* contact within cities where *Haemagogus* are also biting people. Ecological countermeasures, or conservation initiatives, are therefore an effective strategy for primary protection against spillover events; such measures include protecting and restoring wildlife habitat, creating buffer zones, and strategically safeguarding critical habitat for animal feeding, resting, and social aggregation, ultimately strengthening barriers that separate (infected) wildlife and vectors from people (*35*).

Another pillar of epidemic prevention is vaccination. The YFV vaccine is highly efficacious, well studied, and largely available to high risk populations (*11*). Since southeastern Brazil had not experienced a YFV outbreak in nearly a century, most of the region was not considered high risk and was under-vaccinated at the start of the epidemic (*6*). Perhaps more problematic than under-vaccination though, was that in parts of Minas Gerais where YFV vaccinations programs were in place prior to the 2017-18 outbreak, coverage was incomplete and levels of neutralizing antibodies were undetectable across a large proportion of the population (*10, 9*), including in individuals with a documented history of YFV vaccination (*9*). Undetectable levels of neutralizing antibodies indicate waning immunity, which is surprising given the World Health Organization has deemed a single dose of the YFV vaccine confers lifelong immunity, but plausible given some other studies (*40*). This is an important finding as vaccines that do not confer lifelong immunity require different vaccine schedules than those that do and have different implications for long term disease dynamics. Additional issues that should be considered in epidemic prevention planning are potential bottlenecks such as vaccine shortages, cold transport delays, or distribution problems.

The central premise of this study was to investigate the role that drought may have played in triggering an outbreak, building on a growing body of evidence linking precipitation and disease, to ultimately understand how climate and climate change affect infectious disease dynamics. Among the different climate-change related hazards that can affect climate-sensitive infectious diseases, drought is a relatively difficult one to examine. Unlike heatwaves or floods, which are usually distinct events, drought conditions build slowly and can last months to years or more. As a result, robustly linking drought to an epidemic is difficult because of the differing time scales. Despite this challenge, drought has been implicated in other water-borne, vector-borne, and zoonotic disease outbreaks by concentrating humans, animals, and pathogens around limited water sources (*41, 42, 43, 44*). In this study we provide support for two mechanisms for this relationship: 1) drought can cause non-human primate hosts and competent mosquito vectors to move towards human inhabited spaces, and 2) mosquito vectors bite more frequently to avoid desiccation, leading to more opportunities for transmission. Climate change allows droughts to set in quicker and become more intense, so we would expect drought-disease dynamics to become more prominent, yet there are many challenges in developing reliable drought assessments under climate change, making our ability to anticipate such events nontrivial (*45*). On the other hand, the slow build up of drought relative to other events with short lead times like cyclones and floods, provides an opportunity for anticipatory action from health systems as soon as drought conditions start to build. This knowledge could therefore be used to develop early warning systems to better support decision makers and health system planning, both for YFV and potentially other arboviruses. Such systems would be useful beyond Brazil, as other countries like Bolivia, Colombia, Guyana, and Peru face similar deforestation, primate displacement, vaccination, and climate change issues, and have recently seen an uptick in YFV reports (WHO Disease Outbreak News).

## 4 Methods

### 4.1 Model and Parameters

We developed a compartmental model to simulate transmission dynamics for yellow fever virus in Brazil, accounting for key features of the dynamical system. It includes both *Haemagogus* and *Aedes aegypti* mosquitoes as vectors and howler monkeys (genus *Aloutta*), marmosets (genus *Calithrix*), and humans as hosts (Figure 1). Within each species, individuals move between some combination of susceptible, exposed, infectious, and removed (i.e., recovered/dead) states (Figure 1). The model represents transmission in the city and population estimates reflect our understanding of population dynamics around the time of the epidemic: *Aedes aegypti* abundances were approximately 2-3 times lower than in the previous decade due to extensive vector control efforts in response to the Zika pandemic (based on a Rapid Index Survey for *Aedes aegypti* by the Prefeitura Do Rio de Janeiro [accessed 13 Apr 2021]). We estimated *Haemagogus* mosquito abundances as a fraction (55-85%) of the *Aedes aegypti* population because *Haemagogus* mosquitoes typically live in the forest and bite non-human primates in the canopy, only descending and moving to bite people when the forest environment is disturbed or canopy hosts are scarce (*32, 33*). Additionally, we allowed for seasonally varying mosquito carrying capacity, birth rates, and biting rates based on the rainy season. Similarly, we allowed seasonal immigration of howler monkeys and marmosets from the forest to the city as they are highly adapted to fragmented peri-urban habitats (*17, 15, 21, 18*) and many more carcasses were found in and around the city than a typical year (*7*).

The model tracks the number of individuals in the following epidemiological compartments: susceptible (S), exposed (E, where applicable), infectious (I), and recovered/dead (R, where applicable). The compartment subscripts refer to the population group: howler monkeys (p), marmosets (c), humans (h), *Haemagogus* mosquitoes (Hg), and *Aedes aegypti* mosquitoes (aa). For each population group x, the total population size is given by the sum of its compartments:

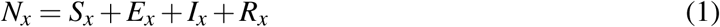

We describe the set of ordinary differential equations below with associated parameter definitions and values in Table 2.

**Table 2:** Model parameter symbols, definitions, values, and sources. Numbers in brackets indicate low and high estimates used for seasonal forcing and/or in the sensitivity analysis.

| Parameter | Definition | Value | Source |
| --- | --- | --- | --- |
| $\alpha$ | Mosquito biting rate (*Haemagogus* and *Aedes*) | 0.70 [0.30, 1.00] | (48, 49, 50) |
| pMI<sub>1</sub> | Probability of *Haemagogus* infection from infectious monkey | 0.20 [1.00] | (51) |
| pMI<sub>2</sub> | Probability of *Haemagogus* infection from infectious human | 0.25 [0.50] | (16) |
| pMI<sub>3</sub> | Probability of *Aedes* infection from infectious human | 0.25 [0.50] | (50) |
| pMI<sub>4</sub> | Probability of *Haemagogus* infection from infectious marmoset | 0.125 [0.25] | (16) |
| $b$ | Probability mosquito becomes infectious after infection | 0.50 | (50) |
| PDR<sub>hg</sub> | YFV development rate in *Haemagogus* | 8.3e−2 | (52, 53) |
| PDR<sub>aa</sub> | YFV development rate in *Aedes* | 7.7e−2 | (54, 55, 56) |
| $\mu_{hg}$ | *Haemagogus* birth/mortality rate | 0.05 [3.70e-02, 1.43e-01] | (57, 58, 59) |
| $\mu_{aa}$ | *Aedes* birth/mortality rate | 0.05 [2.86e-02, 0.10] | (50, 60) |
| $\mu_p$ | Primate birth/mortality rate | 1.6e−4 | (61) |
| $\mu_c$ | Marmoset birth/mortality rate | 4.6e−4 | (62) |
| $\mu_h$ | Human birth/mortality rate | 3.7e−5 | (World Bank) |
| $\mu_{v1}$ | Mortality rate of primates with YFV | 0.50 [0.20, 0.80] | (16) |
| $\mu_{v2}$ | Mortality rate of marmosets with YFV | 0.01 | (16) |
| $\mu_{v3}$ | Mortality rate of humans with YFV | 0.02 | (55) |
| $\gamma_p$ | Primate recovery rate (1/infectious period) | 1.7e−1 | (51, 63, 64) |
| $\gamma_c$ | Marmoset recovery rate (1/infectious period) | 0.25 | (65) |
| $\gamma_h$ | Human recovery rate (1/infectious period) | 2.9e−1 | (51, 50) |
| $\delta_h$ | Human incubation rate (1/incubation period) | 0.23 | (55, 66, 56) |
| $K$ | Mosquito carrying capacity | [4.0e5, 7.0e6] | (this study) |
| $V$ | Vaccination rate (per day) | [0.00, 6.7e−4] | (67, 68) |
| $m_1$ | Movement of howlers from forest to city (#/month) | 100 [5, 200] | (this study) |
| $m_2$ | Movement of marmosets from forest to city (#/month) | 75 [10, 150] | (this study) |
| $m_3$ | Movement of *Haemagogus* mosquitoes from forest to city (#/month) | 10 [1, 100] | (this study) |

**Howler monkeys**

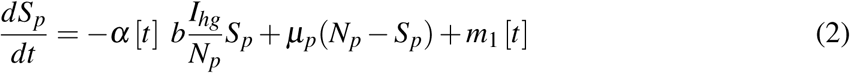

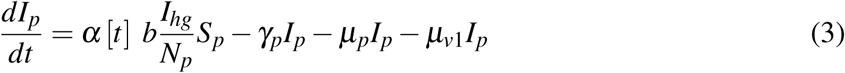

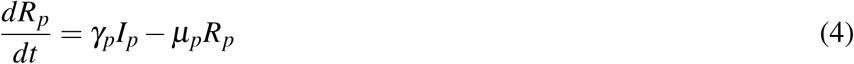

**Marmosets**

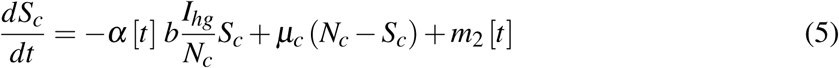

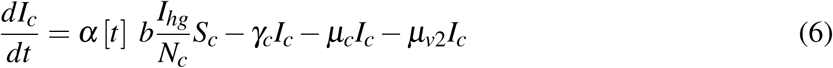

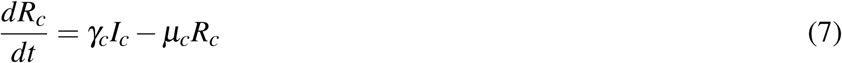

**Humans**

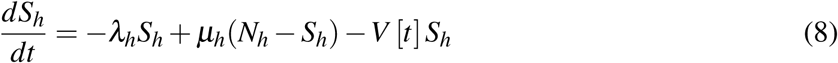

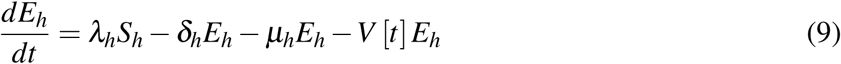

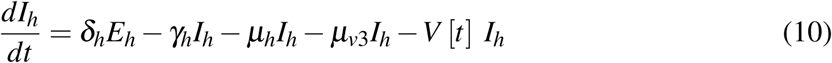

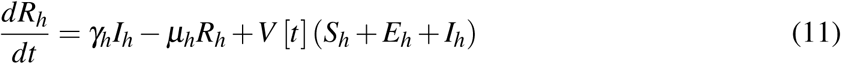

where:

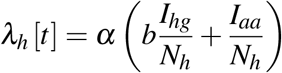

**Haemagogus mosquitoes**

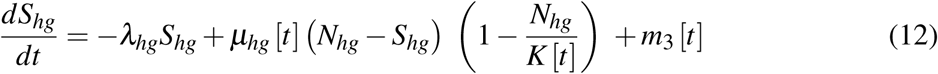

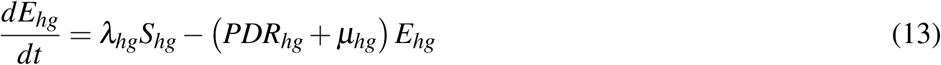

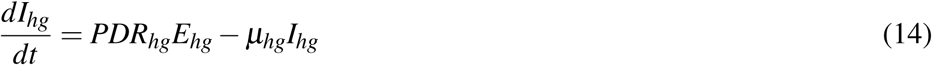

where:

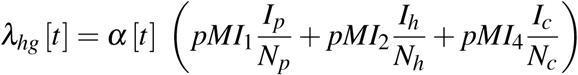

**Aedes aegypti mosquitoes**

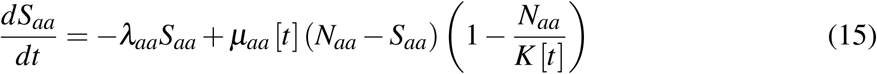

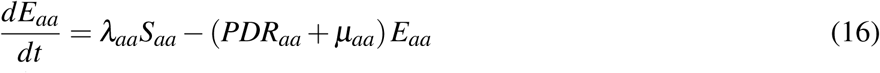

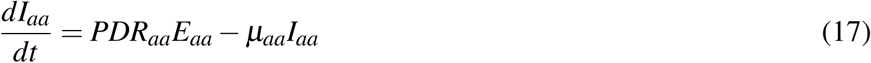

Carrying capacity

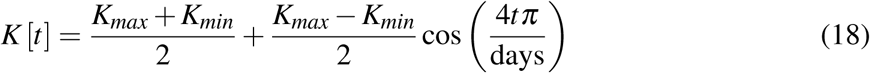

### 4.2 Simulations

To assess the role of rainfall seasonality in driving the YFV epidemic, we simulated disease transmission with four models with different seasonal drivers. The full model included seasonally-driven carrying capacity, mosquito biting rates, and movement (i.e., immigration) of forest species to the city. Carrying capacity is commonly treated as seasonal (e.g., (*46*)), whereas the pervasiveness and importance of seasonally driven biting rates and movement are less well understood. Thus, the three other models tested the importance of these factors in nested versions of the full model: model two included seasonally-driven movement of non-human primates and *Haemagogus* mosquitoes but held mosquito biting rate constant; model three included seasonally-driven mosquito biting but limited movement of forest species; and model four excluded both seasonally-driven movement of forest species and mosquito biting.

Since some parameter values are less certain than others, we assessed the sensitivity of the model outcomes to the most uncertain parameter values. In the sensitivity analysis, we evaluated the impact of increasing and decreasing three highly uncertain parameters resulting in five model specifications. The five models used the full model but replaced the following parameter values: models 1-2) low/high YFV-induced monkey mortality rates; model 3) high probability of mosquito infection with YFV (simultaneously increasing the four pMI values); and models 4-5) low/high movement rate of forest species to the city.

To assess the extent to which different interventions could have prevented the second wave of the YFV epidemic and beyond, we generated four model specifications representing 1) vector control (i.e., reducing mosquito populations by 50% via insecticide spraying in rainy seasons); 2) conservation measures to reduce movement of forest species into urban habitats; 3) earlier deployment of vaccines (three months earlier than actual deployment, corresponding with the initial outbreak, and continuing with early childhood vaccination campaigns); and 4) a combination of all three interventions. We ran the intervention simulations for eight years to assess whether interventions were effective in the short and long term. We additionally ran all four intervention simulations assuming R_0_ were doubled to understand the effectiveness of the interventions if the virus were more transmissible.

For all 17 model specifications (four assessing the role of seasonality, five assessing parameter uncertainty, and eight assessing interventions [four interventions * two R_0_ levels]), we ran each simulation 500 times to account for initial condition uncertainty (i.e., compartment values at the start of the simulation period). We show the distributions for these starting values in table S1.

### 4.3 Data

We obtained monthly confirmed human and non-human primate cases from the Brazilian national health surveillance system, Sistema de Informação de Agravos de Notificação (SINAN). Human cases were confirmed as acute YFV based on serology or polymerase chain reaction (PCR). Non-human primates were tested for YFV by the Division of Zoonoses and Vector-borne Diseases. We used pooled cases of five genera of non-human primates, predominately from the genus *Aloutta* (i.e., howler monkeys), but also composed of *Calithrix* (i.e., marmosets), *Cebus*, *Saimiri*, and *Sapajus*. In the main analysis, we compared simulations with data for the outbreak (November 2016-April 2017; N = 20 months of case counts for both humans and non-human primates). For visu-alization purposes, we show monthly aggregated human cases through 2020 in the intervention plots.

To contextualize the drought preceding the outbreak within historical and future climate conditions, we assessed two metrics of precipitation. We used the Standardized Precipitation Evapotranspiration Index (SPEI) based on weather station data to assess observed conditions between 2007 and 2020 in Minas Gerais, Brazil from (*7*). The index measures water balance, with negative numbers indicating a water deficit. To assess how dry conditions may change in the future in this region, we visualized distributions of historical and future mean monthly evapotranspiration minus precipitation from six models participating in the Coupled Model Intercomparison Project Phase 6 (CMIP6) (*47*): GFDL-CM4, GFDL-ESM4, CESM2-WACCM, CESM2, MPI-ESM1-2-HR, and MPI-ESM1-2-LR. For future projections, we used the SSP2-4.5 scenario, which represents a moderate pathway of greenhouse gas emissions. We estimated return periods by fitting a normal distribution to the precipitation data and using its parameters (mean and standard deviation) to calculate the value corresponding to an exceedance probability of 1/return period. This approach estimates the threshold value that corresponds to a specified frequency of exceedance. To compare the relative magnitude of two exceedance values (e.g., historical vs future 100-yr return period), we calculated the ratio of the larger exceedance value to the smaller one. This ratio quantifies how many times the larger value exceeds the smaller one, providing a sense of their relative difference in terms of magnitude or return period.

### 4.4 Validation

We evaluated how well each model specification (with the exception of intervention scenarios) generated yellow fever virus transmission dynamics by correlating time series of model simulations with observational data. We calculated correlation between monthly predictions and observations using the cross-correlation function in base R. For the non-human primate comparison, we used no time lag, whereas for the human cases we used a one month lag (incorporated as a 30 day lead time of modeled cases). We used a lag for human cases because the observational data was aggregated to monthly case counts, but in reality, cases could have occurred up to 30 days apart. Anecdotally, the infected non-human primate carcasses were found prior to the peak in human cases suggesting a lag of a few weeks, despite the data showing peaks in the two populations in the same month each year. For each model specification, we calculated the correlation between observations (i.e., reported cases/deaths) and median model predictions for humans and non-human primates (howler monkeys only), adjusted based on expected reporting rates (0.7 for non-human primates because mortality rates are high (*9*) and 0.45 for humans based on (*24*)). We determined the best fit model as the model that best characterized both human and non-human primate infection (i.e., highest average correlation across the two categories).

## Acknowledgments

We acknowledge Wenchang Yang for guidance on using CMIP6 data.

## Funding

JMC was supported by the Princeton University Climate and Disease programme with funding from the High Meadows Environmental Institute Grand Challenges and Environmental Studies Strategic Fund and the Joseph and Susan Gatto Foundation. JIR was supported by the National Institutes of Health (NIH K32AI168581).

## Author contributions

Author contribution: JMC and JIR conceived of study and drafted the manuscript. All authors contributed intellectually to methodology, interpretation of results and figures, and manuscript edits.

## Competing interests

“There are no competing interests to declare.”

## Data and materials availability

Data and code are available in the following github repository.

## Supplementary materials

Materials and Methods

Supplementary Text

Figs. S1 to S3

Tables S1 to S4

References (*7–68*)

Movie S1

Data S1

## Sensitivity analysis

The model results were relatively robust to highly uncertain parameters (figure S1, table S2). Increasing and decreasing the YFV-induced mortality rate in monkeys decreased and increased incidence, respectively. In other words, the highest mortality rate resulted in lower incidence, presumably because the monkey population was dying too quickly to amplify transmission. The model was most sensitive to increasing the probability of mosquito infection with YFV: in non-human primates, it resulted in a slightly earlier and higher peak in incidence during the first wave of transmission with an earlier start to the second wave of transmission, while in humans, incidence was substantially higher during both peaks and the second wave occurred earlier than observed. Increasing and decreasing movement rates of forest species increased and decreased incidence, respectively, but also affected the timing of transmission (primarily for low movement rates). The average correlation across all sensitivity analysis simulations was lower than the full model.

---

**Figure S1:**
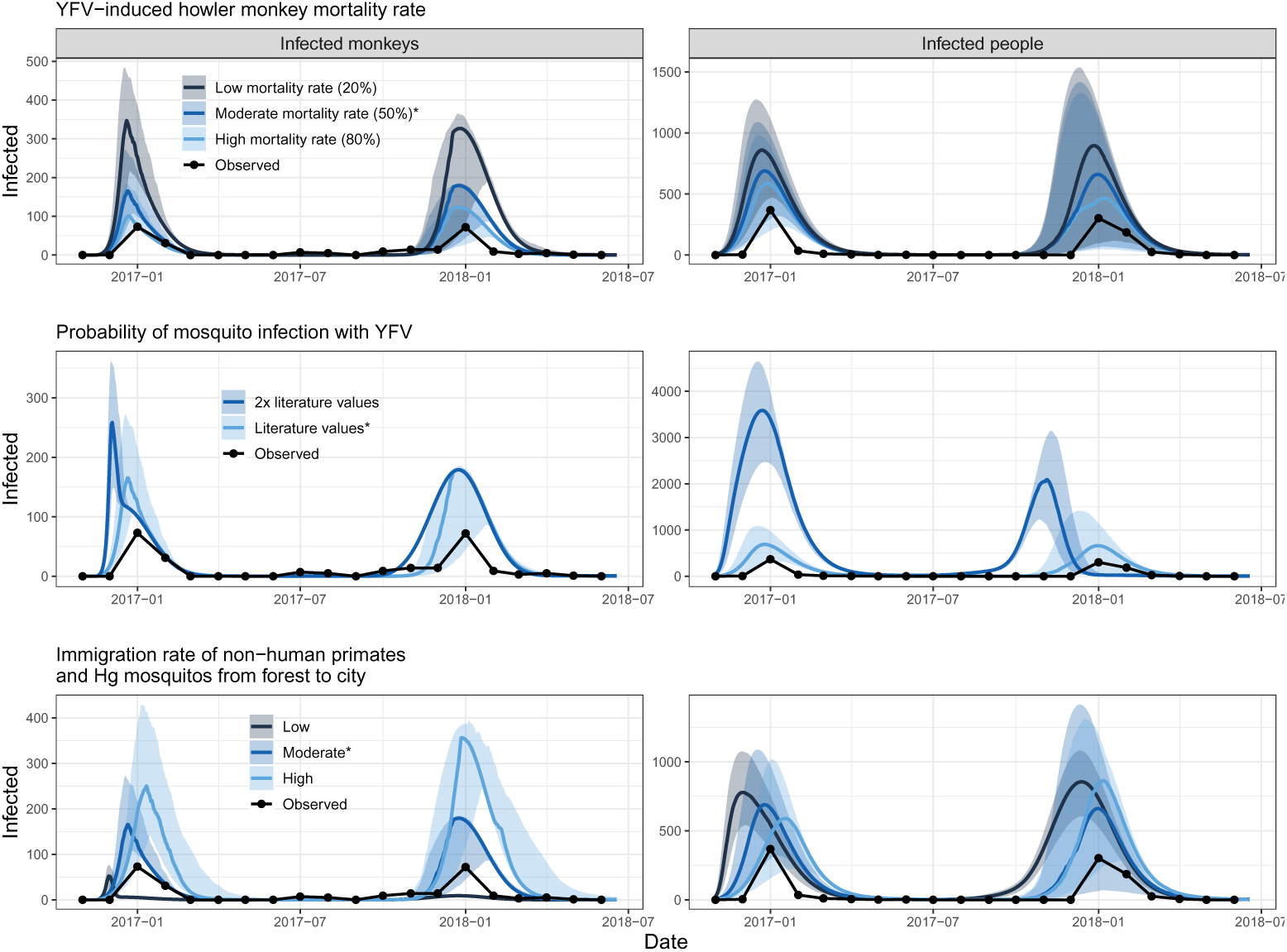
Model results are relatively robust to uncertain parameter values. Infected monkeys (left) and infected people (right). Comparison of model simulations using low and high YFV-induced monkey mortality rates (top), high probability of mosquito infection with YFV (middle), and low and high movement rates for monkeys, marmosets, and *Haemagogus* mosquitoes (bottom). We indicated the values used in the full model with an asterisk and show observations (black points connected with lines) for comparison.

**Table S1:** Distribution functions for generating the 500 initial condition values. Note that the *Haemagogus* mosquito and marmoset populations are proportions of the *Aedes aegypti* and howler monkey populations; thus, the initial conditions for these compartments varied across the simulations accordingly producing variation in initial conditions for five compartments.

| Compartment | Distribution function |
| --- | --- |
| S<sub>aa</sub> | ~Normal(300000, 100000) |
| S<sub>p</sub> | ~Uniform(35000, 10000) |
| R<sub>p</sub> | ~Normal(0.7, 0.1) |

**Table S2:** Correlation (and p-value) between model predictions and observations across nested seasonally-driven models. Modeled human cases were adjusted by a one month lead time. NHP: non-human primates. Note that the’No seasonality’ model still includes a seasonal mosquito carrying capacity. We included the full model in the table for comparison.

| Model | Human cases | NHP cases | Average correlation |
| --- | --- | --- | --- |
| Full model | 0.90 (p <0.001) | 0.90 (p <0.001) | 0.90 |
| Low YFV-monkey induced mortality rate | 0.87 (p <0.001) | 0.91 (p <0.001) | 0.89 |
| High YFV-monkey induced mortality rate | 0.88 (p <0.001) | 0.89 (p <0.001) | 0.89 |
| High probability of mosquito infection with YFV | 0.42 (p <0.05) | 0.60 (p <0.05) | 0.51 |
| Low movement rate of forest species | 0.54 (p <0.01) | 0.09 (p >0.05) | 0.32 |
| High movement rate of forest species | 0.82 (p <0.001) | 0.88 (p <0.001) | 0.85 |

## Notes

https://github.com/jms5151/YFV_Brazil

### Competing Interest Statement

The authors have declared no competing interest.

### Summary of Updates

This version was revised with an updated model and associated simulation outputs. We also added some references and fixed grammatical errors.

